# CryoMV: Structure-Prior-Guided Modeling and Real-Particle Validation of Continuous Conformational Transitions in Cryo-EM

**DOI:** 10.64898/2026.09.21.753206

**Authors:** Fuwei Li, Hao Dong, Shuai Tang, Xinsheng Wang, Chuanyang Zhang, Chenxuan Ji, Renmin Han, Fa Zhang, Xiaohua Wan

## Abstract

Continuous protein conformations are essential for understanding fundamental biological processes and supporting drug discovery. Although cryo-EM can resolve individual states at high resolution, recovering continuous heterogeneity from 2D particle images remains challenging. High noise, motion blur, and limited structural priors make it difficult to accurately generate and validate high-resolution continuous conformations using raw particle data. Here, we introduce cryoMV, a framework that integrates structure-prior-guided modeling with real-particle validation for continuous conformational transitions. CryoMV uses reference density maps to establish structural anchors and motion priors, models candidate transition paths between selected conformations, and transfers the learned representation to raw 2D cryo-EM particle images. Each candidate conformation is subsequently evaluated using the estimated particle poses and contrast transfer functions. Supported conformations are reconstructed through raw particle back-projection and assessed using canonical half-maps and Fourier shell correlation. On EMPIAR-10516 and EMPIAR-10345, cryoMV achieves excellent performance in terms of robustness, verifiability, and reconstruction resolution. By incorporating structure-prior modeling and evidence from the raw particles, cryoMV offers an explicit mechanism for assessing whether generated conformations are supported by experimental data and provides a practical approach to reducing model-induced artifacts in continuous cryo-EM heterogeneity analysis.

## Introduction

Single-particle cryo-electron microscopy (cryo-EM) reconstructs the three-dimensional structure of a protein from a large number of noisy two-dimensional particle projections (Bai, McMullan, and Scheres 2015). Proteins may adopt multiple conformations during imaging, and a single consensus structure cannot capture the full conformational landscape. Such conformational dynamics are closely coupled to molecular function and ligand binding (Henzler-Wildman and Kern 2007), while shifts in conformational ensembles can mediate allosteric regulation (Motlagh et al. 2014); characterizing them can support mechanistic studies and drug design. However, recovering continuous conformational states from cryo-EM data remains difficult because individual particle images have low signal-to-noise ratios and provide only two-dimensional observations with unknown poses and conformations. Reconstructing and validating these states against raw particle data is therefore a central challenge in cryo-EM heterogeneity analysis.

Recovering continuous conformational variability from noisy 2D cryo-EM images remains a challenging inverse problem (Sorzano et al. 2019). Recent advances in 3D generative networks have improved the modeling of continuous heterogeneity. Density generation models like cryo-DRGN2 (Zhong et al. 2021b) encode each particle into a low-dimensional latent space and decode 3D densities. Deformation-based methods such as 3DFlex (Punjani and Fleet 2023) and DynaMight (Schwab et al. 2024) model structural motion via a canonical density and deformation fields. CryoSPARC 3D Variability Analysis (Punjani and Fleet 2021) and RECOVAR (Gilles and Singer 2025) capture conformational variability through low-dimensional covariance analysis and kernel regression. HetSIREN (Herreros et al. 2025) reconstructs continuous conformations using a sinusoidal neural network with decoupled architectures.

Although these methods can analyze conformational heterogeneity in continuous protein motion, their standard workflows typically focus on a single target dataset. They start from one particle stack and learn conformational changes within that stack relative to a consensus 3D structure. Existing algorithms struggle to jointly use multiple particle stacks and their associated 3D structural motion prior networks to learn physically realistic motion processes. As a result, these methods do not explicitly constrain latent-space exploration to transitions between selected reference conformations. More generally, cryo-EM reconstruction pipelines can be affected by algorithmic bias and overfitting; therefore, generated structural changes should be validated against the observed particle images (Sorzano et al. 2022).

To address the questions of which structural changes should be explored and which motions are valid and reliable, we propose cryoMV, a continuous heterogeneity reconstruction framework that combines structure-prior-guided mod-eling, real-particle validation, and evidence-driven transfer. First, the Motion Prior Module establishes direct motion mappings between different 3D conformational states and infers both flexible and consensus regions. Second, the Fourier Generation Module learns motion states of different 3D conformations, encoding each conformation and its motion mappings as points and lines in the latent space. The inferred flexible regions constrain generated motions to these areas, thereby improving motion realism. This design enhances conformational separation and provides an initial mapping of motion trajectories. Third, the Particle-Evidence Transfer Module verifies the authenticity of generated conformations through real 2D particle classification and reconstruction. The reconstructed particle states are then used as inputs to the neural network, and transfer learning is applied to establish a mapping between 3D structural changes and 2D particle images. Finally, the Particle-Evidence Validation Module re-decodes candidate structures and revalidates the updated conformational motion states, implementing an evidence-feedback loop of “generation–validation–updated generation–re-validation.” We evaluated cryoMV on two publicly available cryo-EM datasets, EMPIAR-10345(Campbell et al. 2020) and EMPIAR-10516(Melero et al. 2020); the former contains multiple small movements, while the latter contains localized large-amplitude movements. Under metrics for raw particle verifiability and robustness, cryoMV achieved state-of-the-art performance, demonstrating its ability to generate reliable dynamic conformations. In terms of 3D reconstruction resolution metrics, cryoMV achieved results close to the current best and near-best on both datasets.

## Related Work

### Traditional particle classification and 3D reconstruction methods

Traditional cryo-EM heterogeneity analysis groups particles into discrete classes, iteratively assigning particles and updating class densities. RELION (Scheres 2012) applies Bayesian MAP estimation for classification and reconstruction, while cryoSPARC (Punjani et al. 2017) uses stochastic optimization for rapid 3D classification. These methods jointly estimate particle classes, pose, and corresponding 3D structure from many noisy 2D particle images with unknown orientations, forming the basis of cryo-EM classification and reconstruction. However, standard discrete classification usually requires the number of classes to be specified in advance, producing a finite set of class structures that cannot easily describe the motion between different conformations.

### Covariance-based statistical heterogeneity analysis methods

Covariance-based statistical heterogeneity analysis methods estimate low-dimensional covariance in 3D density to extract main modes of variation. CryoSPARC 3D Variability Analysis (Punjani and Fleet 2021) uses probabilistic PCA for this purpose, and RECOVAR (Gilles and Singer 2025) combines covariance estimation, probabilistic coordinates, kernel regression, and uncertainty analysis to reconstruct structures and analyze transitions. These methods are statistically interpretable and do not require generative networks, but low SNR or unreliable covariance may mix real motions with noise, leading to inaccurate results.

### Latent-variable implicit density generation methods

To further capture nonlinear conformational changes, density generative methods encode particles in latent space and decode 3D densities, capturing nonlinear conformational changes. CryoDRGN (Zhong et al. 2021a) and CryoDRGN2 (Zhong et al. 2021b) use coordinate-conditional networks and pose estimation to model continuous structural distributions, enabling latent space sampling. However, they are usually trained on single datasets and provide limited insight into multi-conformational transitions.

### Deformation-field–based conformational modeling methods

Unlike density generation methods, deformation-field approaches model conformational changes as continuous transformations of a reference, constrained by structural or geometric priors. Zernike3D (Herreros et al. 2023) uses 3D Zernike polynomials for interpretable deformations. 3DFlex (Punjani and Fleet 2023) and DynaMight (Schwab et al. 2024) model deformations while preserving geometry. CryoSTAR (Li et al. 2024)adds atomic models and regularization. Reference structures and geometric constraints improve motion plausibility, but inferring continuous 3D deformations from noisy 2D images remains challenging, especially in regions with weak motion or limited evidence.

### Decoupled heterogeneity reconstruction methods

Some methods improve heterogeneity reconstruction by decoupling conformation from imaging factors. OPUS-DSD (Luo et al. 2023) uses a 3D convolutional encoder–decoder for structural decoupling, while HetSIREN (Herreros et al. 2025) employs a sine representation and supernet for realspace reconstruction, enhancing stability by decoupling conformation, pose, and CTF. These approaches reduce confounding factors and improve representation learning but often incur high computational costs and may require aggressive downsampling, leading to structural detail loss.

## Method

As shown in Fig. 1, the cryoMV framework has four modules. The Motion Prior Module derives direct motion trajectories from reference conformations and identifies local motion regions. The Fourier Generation Module uses multiple reference density maps and their direct motion trajectories to learn structural variation prior and constrain these regions. The Particle-Evidence Transfer Module checks Fourier-domain consistency between the generated map and the original 2D particles and updates the network. The Particle-Evidence Validation Module then re-evaluates the updated map and uses back-projection to recover a particle-consistent conformation. Through this process, cryoMV can identify protein motions that are supported by the 2D evidence.

**Figure 1.**
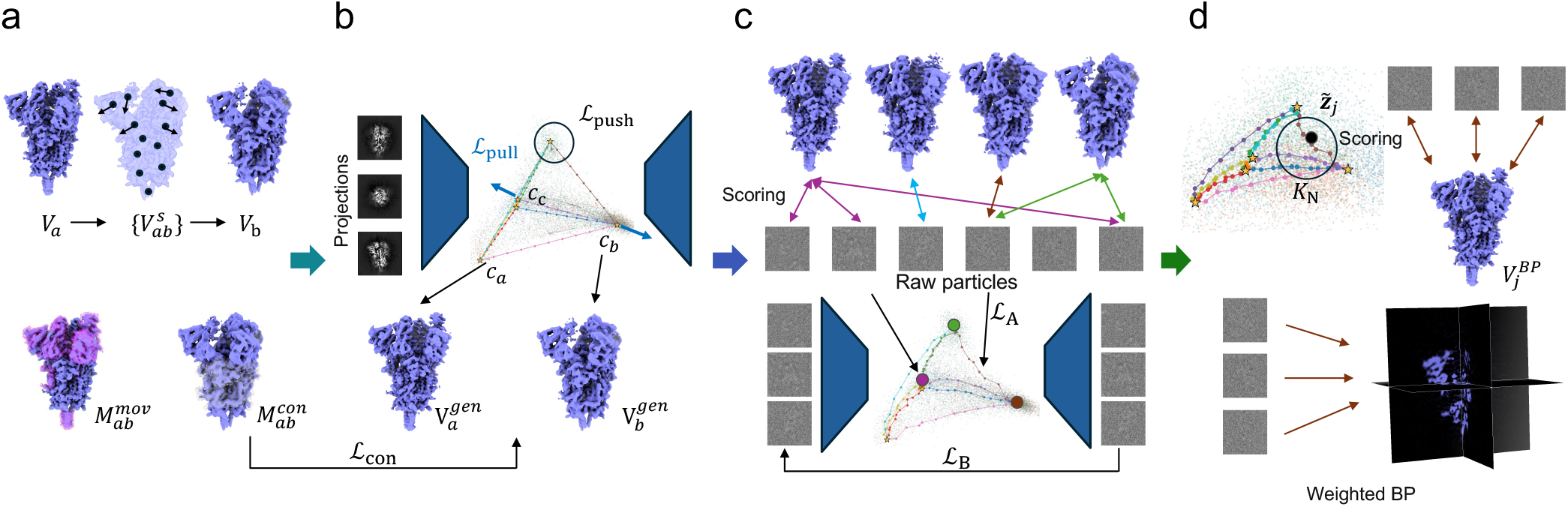
Overview of cryoMV. (a) The Motion Prior Module derives direct transition trajectories and region masks from the reference conformations. (b) The Fourier Generation Module learns the latent space from simulated projections and direct transition trajectories between reference states. (c) The Particle-Evidence Transfer Module scores candidate conformations against raw particles and transfers the latent representation toward the particle domain. (d) The Particle-Evidence Validation Module groups nearby particles in latent space and reconstructs the map to compute weights. At last, weighted back-projection is used to obtain the final structure.

### Motion Prior Module

The Motion Prior Module extracts two prior knowledge for each pair of conformations: the direct motion trajectory, and mask (Fig. 1 a).

To remove structural noise, for each reference conformation pair (*V*_*a*_, *V*_*b*_) with density thresholds *q*_*a*_ and *q*_*b*_, we remove low-density voxels and normalize the remaining density:

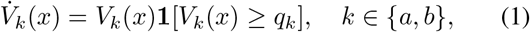

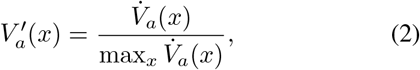

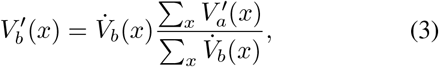

where 1[*·*] denotes binarization. Next, the Motion Prior Module converts the matched initial density 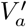 into a weighted point cloud, where the spatial coordinates of the particles are 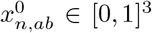, and the density weights are denoted as 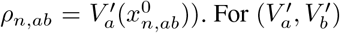, the model independently trains a velocity network *v*_*ψ,ab*_. Throughout the process, point positions are updated via *K* steps of explicit Euler integration (Fan et al. 2024):

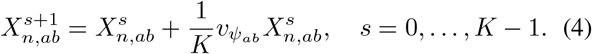

For each point, use trilinear interpolation *S* to map the weight back to the density map:

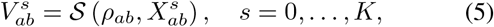

where *v*_*ψ*_ is optimized using the Wasserstein loss (Cuturi 2013):

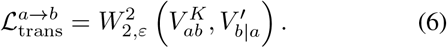

After training, we can obtain the complete point states 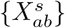 and the corresponding direct motion trajectories 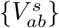.

However, 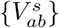 may contain global translations and rotations in (*V*_*a*_, *V*_*b*_). To highlight local conformational differences, we find an affine transformation 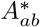 from the initial position 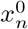 to the final position 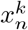, weighted by density:

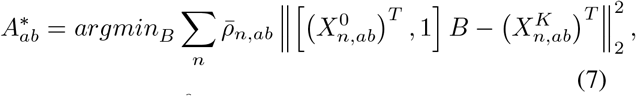

where 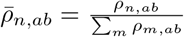. The distance after an affine transformation *d*_*n,ab*_ can be expressed as:

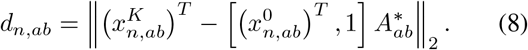

This yields the motion region mask *M*_*ab*_:

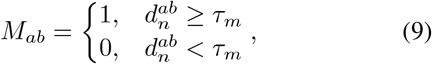

where *τ*_*m*_ is a hyperparameter that determines the size of the flexible region. Next, 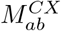 is obtained by segmenting the density map using ChimeraX (Pettersen et al. 2021), and the motion region is defined as:

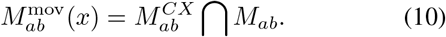

The consensus region can be expressed as:

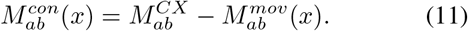

Thus, we have obtained the consensus region mask 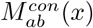 and the direct motion trajectories 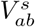 of the reference conformation pair (*V*_*a*_, *V*_*b*_).

### Fourier Generation Module

The Fourier Generation Module learns protein structures and motions from simulated images *Y*_*i*_ made from reference density *V* maps and their motions 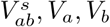 ∈ *V* (Fig. 1 b). The mask 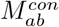 will keep the consensus structure from moving.

For each projection image *Y*_*i*_, it is first transformed into Fourier space *Ŷ*_*i*_ and fed into the encoder to predict the mean *µ*_*i*_ and log-variance 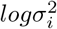 of the Gaussian posterior. Its latent variables are obtained through reparameterization(Kingma and Welling 2013)

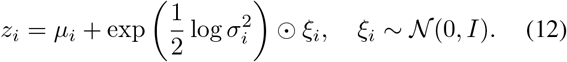

The decoder *G*_*θ*_(*z, k*) predicts the Fourier representation of the density at position *k* for a given latent variable *z*_*i*_:

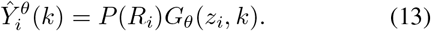

In particular, *R*_*i*_ represents the projection pose of the simulated image *Y*_*i*_, and *P* (*·*) denotes the Fourier slice operator. Therefore, the Fourier projection reconstruction loss is

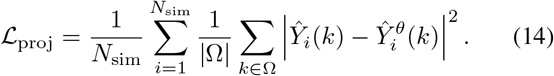

Here, Ω denotes the actual Fourier region used for training, and *N*_*sim*_ is the number of simulated projections. To make sure the latent space clearly matches the known reference conformations, we assign cluster centers (*c*_*a*_, *c*_*b*_) in the latent space to each reference conformation pair 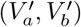. Then, we use a center separation loss to push different reference conformations as far apart as possible:

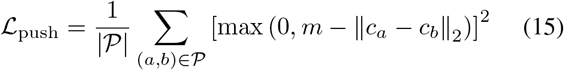

Here, *P* denotes all distinct pairs of reference centers, and *m* is the minimum distance between them. For its direct motion trajectory 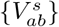, we constrain the corresponding cluster centers as follows:

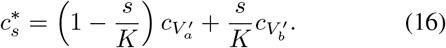

Therefore, 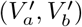 are represented as discrete anchor points in the latent space, while 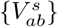 is encoded as the lines connecting the anchor points. The projection *Y*_*i*_ encodes the mean, which is constrained to the target position by the following equation:

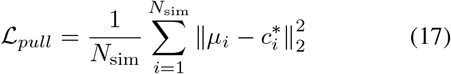

Therefore, the latent space central loss can be written as

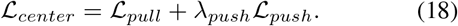

We then impose constraints on the decoding structure by using 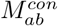 to mark regions where changes are not permitted. For the densities *G*_*θ*_(*c*_*b*_, *k*) and *G*_*θ*_(*c*_*a*_, *k*) decoded by the generative network at the reference centers *c*_*a*_ and *c*_*b*_, the loss for the fixed regions is defined as:

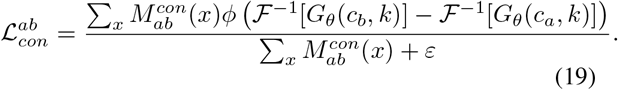

Here, 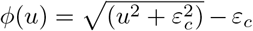 is the Charbonnier (Charbonnier et al. 1994) penalty function. The training objective is:

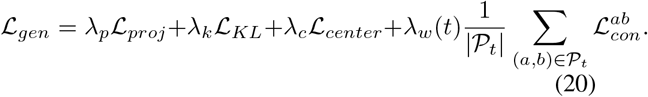

where *λ*_*p*_, *λ*_*k*_, and *λ*_*c*_ are hyperparameters; *L*_*KL*_ is the standard KL loss (Kingma and Welling 2013); and *λ*_*w*_(*t*) is the warmup function at training step *t*. After completing the first training phase, we validate the generated results and use the validation data to further optimize the Fourier Generation network.

### Particle-Evidence Transfer Module

The Particle-Evidence Transfer Module converts the trained latent space into a 3D density map and finds high-confidence structures and their related particle stacks. It uses transfer learning to move the learned conformational representations from the reference projection domain to the original particle domain (Fig. 1 c). Specifically, cryoMV performs elliptical sampling on each pair of reference structures (*c*_*a*_, *c*_*b*_) within the latent space, and the candidate latent variables are represented as:

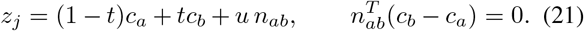

Here, *t* controls the position in (*c*_*a*_, *c*_*b*_), and *u* controls the lateral disturbance. For a candidate coordinate *z*_*j*_, the decoder generates the corresponding 3D density:

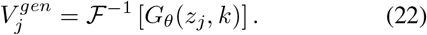

Next, the Particle-Evidence Transfer Module calculates the Fourier-domain consistency between each 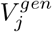and each original particle *I*_*i*_ under pose *R*_*i*_ and fixed CTF conditions:

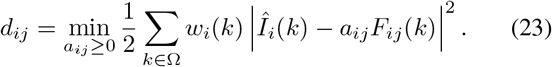

The *a*_*ij*_ adjusts for overall amplitude differences, *w*_*i*_(*k*) is noise-related frequency weighting, *F*_*ij*_(*k*) is the predicted spectrum of 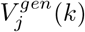 (Singer and Sigworth 2020):

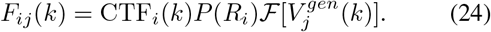

We use the error of the blank projection 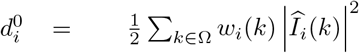 as a reference to get evidence score:

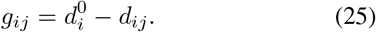

For each generated 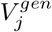, cryoMV sorts the particles *I*_*i*_ according to *g*_*ij*_ and takes the mean of the top *K* values as the score for 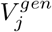. For each pair (*V*_*a*_, *V*_*b*_), it retains the 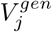 with the higher score and its corresponding *z*_*j*_, ultimately yielding *J* credible structures. For each 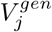, the particles are again sorted according to *g*_*ij*_ to obtain *r*_*ij*_, which is then used to calculate the particle support weights:

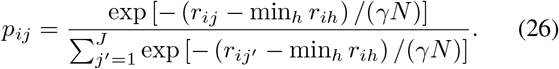

Here, *γ* is a hyperparameter. A smaller *γ* makes the weights focus more on the top *V*_*j*_. *N* is the true number of particles. Therefore, the evidence-weighted latent space center of *I*_*i*_ is

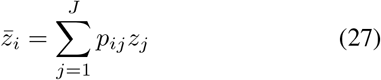

After determining the particle’s position in the latent space, the Particle-Evidence Transfer Module transfers the original particle features to the existing conformational latent space in two stages. In the first stage, the Fourier decoder is frozen, and only the encoder is updated. Given that high-noise particles typically cannot be reliably localized to a unique latent point, the model does not directly set the encoder mean equal to 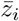, but instead samples the target *u*_*i*_ within a local latent sphere centered at 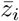. The radius of this sphere is determined by the average distance between adjacent candidate nodes. The loss function for the first stage is

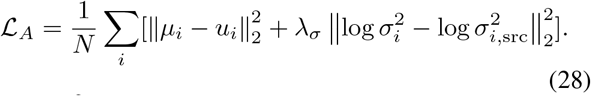

The 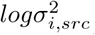 is generated by the frozen source encoder.

The second stage further adapts the Fourier decoder. To limit latent space drift at the start of joint updating, the model uses the frozen first-stage encoder as teacher:

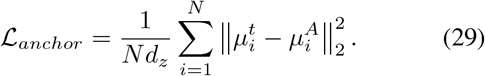

Here, 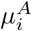 is the mean of the latent variables output by the encoder frozen in the first stage for particle *I*_*i*_, and 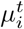 is the output of the current encoder for the same particle. Then the CTF-aware reconstruction loss *L*_*CTF*_ can update the encoder and decoder jointly:

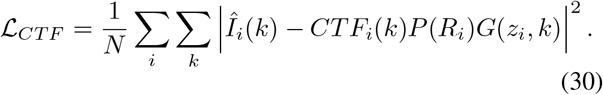

Thus, the joint training objective is

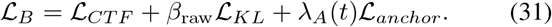

The *L*_KL_ is the standard KL loss, and *λ*_*A*_(*t*) is a decay schedule function with respect to training time *t, L*_*anchor*_ is used to constrain latent space drift during the joint training phase.

### Particle-Evidence Validation Module

This module verifies the generated conformations and refines the reconstruction results by combining prior structural information with evidence from the original particles (Fig. 1 d). First, after the Particle-Evidence Transfer Module, the latent space has changed. We need to align the cluster centers of conformations 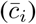 from the reference space with their new positions 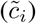 in the updated latent space:

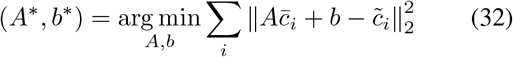

Then, each coordinate *z*_*j*_ in the new latent space is mapped using this transformation:

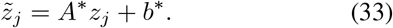

Based on these updated latent coordinates, the 3D model is reconstructed as:

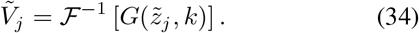

Next, we use the 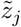 positions to find the K-nearest neighbors *N*_*K*_(*j*) in latent space. These neighbors are used to reconstruct the density map and compute validation scores for the particles. Based on these scores, back-projection is performed on the original particles

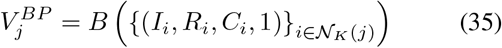

Here, *B* denotes back-projection. The particles with the highest validation scores form the set *T*_*j*_. Their scores are normalized to [0, 1] to get the back-projection weights *ω*_*ij*_, and the final structure is reconstructed using weighted back-projection:

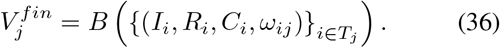

Through this closed-loop process of “generation—validation,” cryoMV integrates the conformational prior from the reference with evidence from observed particles, enabling reliable identification of dynamic conformations.

## Experiments

This section evaluates cryoMV from 4 aspects: (1) the reslution of generated and reconstructed 3D conformations; (2) the verifiability of generated conformations and motions; (3) the model’s robustness in conformation identification and reconstruction; and (4) the contribution of each module within cryoMV. The visualization of the motion process are shown in Supplementary Material.

## Implementation Details

### Experimental Setup

We evaluate EMPIAR-10516, which contains large localized motion, and EMPIAR-10345, which contains multiple subtle motions. CryoMV is compared with CryoDRGN2, OPUS-DSD, DynaMight, and RECOVAR using identical particles, masks, poses, and CTFs, together with a unified center-matching, particle-selection, and back-projection protocol. We evaluate 105 states from five centers on EMPIAR-10516 and 64 states from four centers on EMPIAR-10345.

### Training Configuration

For training CryoMV, we used a three-layer velocity field network to model motion priors, with a hidden layer width of 100, and discretized the conformational motion into 10 transition frames. The Fourier generative model runs on a 256^3^ grid with 8 latent dimensions and a hidden layer width of 1,024. It is trained for 10 epochs using the AdamW optimizer with a learning rate of 1*×*10^*−*4^ and a global batch size of 20. Particle evidence transfer uses a learning rate of 1*×*10^*−*5^: In Phase A, the decoder is frozen and the encoder is trained for 10 epochs; in Phase B, all particles are used for joint training for 10 epochs, with a global batch size of 32.

### Evaluation Metrics

We report GS-FSC (Scheres and Chen 2012; Rosenthal and Henderson 2003) for the resolution of particle-supported reconstructions, Map2Map FSC and masked SSIM (Zeng and Wang 2012) for the consistency between generated maps and particle back-projections, and Mixed% for the contaminant fraction among the top 50,000 selected particles. Lower FSC and Mixed% and higher SSIM indicate better performance. All consistency metrics are averaged over matched conformational centers within the motion mask. For detailed metrics explanation, see the Supplementary Material.

## Main Results

### Reconstruction Resolution

As shown in Table 1, cryoMV achieved the best GS-FSC resolution on EMPIAR-10516 and ranked third in Map-to-Map FSC, just behind cryo-DRGN2 and RECOVAR. However, Fig. 2(a) shows that cry-oDRGN2 and RECOVAR’s results contain substantial noise and artifacts, suggesting overfitting, while cryoMV produces smoother structures that are consistent with particle back-projection. On EMPIAR-10345, cryoDRGN2 reached the highest GS-FSC resolution, with cryoMV trailing by only 0.191 Å. Both methods achieved the best Map-to-Map FSC of 4.623 Å. Additional consistency analyses further demonstrate that cryoMV outperforms other methods and avoids overfitting artifacts. Overall, cryoMV delivers near state-of-the-art resolution in particle reconstruction and leads in consistency and reliability of the generated conformations.

**Table 1:** Reconstruction resolution on EMPIAR-10516 and EMPIAR-10345. Lower is better.

|  | Method | GS-FSC (Å) ↓ | Map2Map FSC (Å) ↓ |
| --- | --- | --- | --- |
| EMPIAR-10516 | CryoMV | **4.435** | 4.027 |
|  | CryoDRGN2 | 4.476 | **2.487** |
|  | OPUS-DSD | 4.723 | 7.255 |
|  | DynaMight | 4.525 | 7.999 |
|  | RECOVAR | 4.659 | 3.741 |
| EMPIAR-10345 | CryoMV | 5.246 | **4.623** |
|  | CryoDRGN2 | **5.055** | **4.623** |
|  | OPUS-DSD | 5.206 | 18.445 |
|  | DynaMight | 5.172 | 5.368 |
|  | RECOVAR | 6.390 | **4.623** |

**Figure 2.**
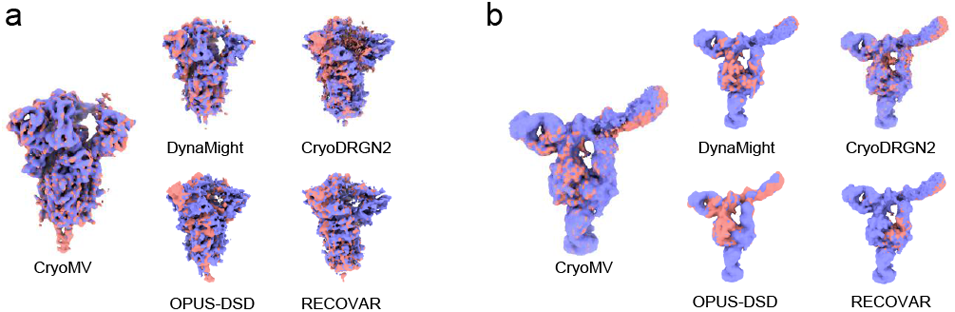
Comparison of generated map and back-projection map. Purple represents the BP results, and red represents the generated results; the more similar they are, the better. (a) Comparison on EMPIAR-10516. (b) Comparison on EMPIAR-10345.

### Generation Verifiability

As shown in Tab. 2, cryoMV leads in multiple generative consistency metrics on both datasets. On EMPIAR-10516, it achieved the highest C-SSIM and P-SSIM scores, with only C-FSC and P-FSC scores slightly below those of cryoDRGN2. However, Fig. 2(a) shows that cryoDRGN2’s lower FSC values are accompanied by high-frequency noise and artifacts, indicating overfitting and reducing reliability. In contrast, cryoMV’s generated structures are smoother and more consistent with particle back-projections, without obvious artifacts. For EMPIAR-10345, which features subtle local motions, most methods show discrepancies between generated and back-projected structures. Among them, cryoMV and RECOVAR deliver relatively stable results.

**Table 2:** Structural consistency on EMPIAR-10516 and EMPIAR-10345. C and P denote center-level and pair-level evaluation, respectively. Higher SSIM and lower FSC resolution indicate better performance.

| Method | EMPIAR-10516 |  |  |  | EMPIAR-10345 |  |  |  |
| --- | --- | --- | --- | --- | --- | --- | --- | --- |
| | C-SSIM ↑ | P-SSIM ↑ | C-FSC ($\text{\AA}$) ↓ | P-FSC ($\text{\AA}$) ↓ | C-SSIM ↑ | P-SSIM ↑ | C-FSC ($\text{\AA}$) ↓ | P-FSC ($\text{\AA}$) ↓ |
| CryoMV | **0.799** | **0.804** | 4.016 | 4.027 | **0.867** | **0.889** | **4.623** | **4.623** |
| CryoDRGN2 | 0.619 | 0.623 | **2.516** | **2.485** | 0.832 | 0.856 | **4.623** | **4.623** |
| OPUS-DSD | 0.702 | 0.714 | 7.258 | 7.255 | 0.725 | 0.727 | 15.191 | 18.465 |
| DynaMight | 0.468 | 0.474 | 8.052 | 7.998 | 0.857 | 0.877 | 5.416 | 5.367 |
| RECOVAR | 0.619 | 0.613 | 3.747 | 3.741 | 0.812 | 0.811 | **4.623** | **4.623** |

We compared the motions generated by each method with the ground truth. In Fig. 3(a), cryoMV accurately reproduced both significant local motions, while cryoDRGN2 detected motion in incorrect regions, DynaMight identified three regions, RECOVAR detected one, and OPUS-DSD detected none. In the PCA analysis, cryoMV clearly matched the ground truth cluster centers and showed interpretable motion involving many particles, whereas other methods showed only central clustering with limited interpretability. In Fig. 3(b), cryoMV again captured both small-amplitude local motions. RECOVAR and OPUS-DSD also detected these motions, though OPUS-DSD’s results were less pronounced. DynaMight detected only one, with incorrect direction. In the PCA plots, only cryoMV consistently demonstrated clear conformational separation and interpretable motion, while other methods produced a single cluster lacking interpretability.

**Figure 3.**
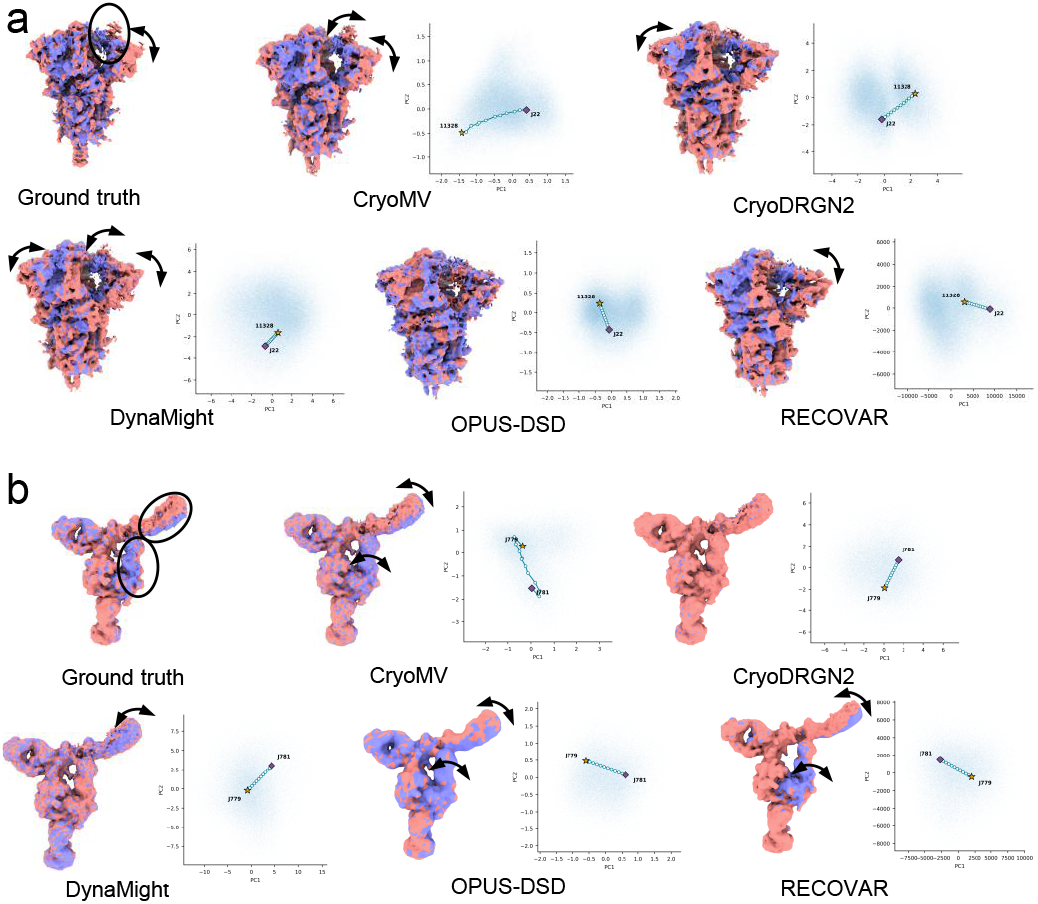
Comparison of generated motions with ground truth. Purple and red maps represent different conformations; arrows and circles highlight the regions and directions of motion. (a) Results on EMPIAR-10516. (b) Results on EMPIAR-10345.

Overall, quantitative metrics, visualizations, and motion analyses demonstrate that cryoMV achieves state-of-the-art performance in conformational consistency and verifiability.

### Robustness

To assess algorithm robustness, we added contaminant particles (noise and particles from other cryo-EM datasets) to the EMPIAR-10516 particle stacks at proportions of 10% and 30% of the particles. Because not all particles are used in 105 reconstruction, the contamination ratio in reconstruction stacks may exceed 30%.

As shown in Tab. 3, cryoMV outperformed all other methods on every metric under both 10% and 30% particle contamination. At 10% contamination, cryoMV had the best GS-FSC, M2M, and C-SSIM scores, and selected far fewer contaminant particles than other methods. Even at 30% contamination, its performance and contaminant exclusion remained strong, while other methods degraded significantly. These results highlight cryoMV’s superior robustness and resistance to contamination.

**Table 3:** Robustness under 10% and 30% particle contamination (on EMPIAR-10516). M2M and C-FSC denote Map2Map FSC and center-level Map2Map FSC, respectively. FSC values are reported in Å. Higher C-SSIM and lower values for the other metrics indicate better performance.

| Method | 10% Contamination |  |  |  |  | 30% Contamination |  |  |  |  |
| --- | --- | --- | --- | --- | --- | --- | --- | --- | --- | --- |
| | GS-FSC ↓ | M2M ↓ | C-SSIM ↑ | C-FSC ↓ | Mixed ↓ | GS-FSC ↓ | M2M ↓ | C-SSIM ↑ | C-FSC ↓ | Mixed ↓ |
| CryoMV | **4.446** | **3.831** | **0.733** | **3.750** | **2.464** | **4.440** | **3.956** | **0.715** | **3.802** | **7.806** |
| CryoDRGN2 | 4.598 | 4.197 | 0.478 | 4.197 | 5.750 | 4.599 | 4.134 | 0.469 | 4.134 | 20.842 |
| OPUS-DSD | 4.896 | 18.218 | 0.509 | 18.218 | 5.832 | 4.796 | 7.456 | 0.503 | 7.456 | 22.256 |
| DynaMight | 4.576 | 7.383 | 0.388 | 7.525 | 2.572 | 4.704 | 7.595 | 0.352 | 7.522 | 17.934 |
| RECOVAR | 4.696 | 15.385 | 0.508 | 20.993 | 8.748 | 4.656 | 6.248 | 0.532 | 5.549 | 32.803 |

### Ablation Experiment

Table 4 presents ablation results on EMPIAR-10516, where ET (Evidence Transfer) and MP (Motion Prior) are key modules. Removing ET caused a marked decline in all metrics. For example, C-SSIM and P-SSIM dropped from 0.799 and 0.804 to 0.482 and 0.484, respectively, while all FSC values increased. These results demonstrate that evidence transfer is crucial for integrating complementary information and improving reconstruction quality. Removing MP caused a sharp decrease in P-SSIM and notable increases in P-FSC, GS-FSC, and M2M FSC, indicating that motion priors are essential for capturing particle motion and conformational correspondences. Overall, the complete cryoMV model achieved the best or near-best results across all metrics, confirming that ET and MP provide complementary benefits in terms of conformation consistency, verifiability, and reconstruction resolution.

**Table 4:** Ablation results on EMPIAR-10516. ET and MP denote Evidence Transfer and Motion Prior, respectively.

| Metric | CryoMV | w/o ET | w/o MP |
| --- | --- | --- | --- |
| C-SSIM ↑ | 0.799 | 0.482 | **0.848** |
| P-SSIM ↑ | **0.804** | 0.484 | 0.425 |
| C-FSC ($\text{\AA}$) ↓ | 4.016 | 4.866 | **3.936** |
| P-FSC ($\text{\AA}$) ↓ | **4.027** | 4.808 | 5.414 |
| GS-FSC ($\text{\AA}$) ↓ | **4.435** | 4.640 | 8.838 |
| M2M FSC ($\text{\AA}$) ↓ | **4.027** | 4.812 | 5.344 |

## Conclusion

This paper introduces cryoMV, a framework for continuous conformation reconstruction that integrates reference structure priors with raw particle data. CryoMV extracts motion trajectories and local motion regions from known conformations, generates candidate structures in the Fourier domain, and validates them using particle matching, latent space migration, and weighted back-projection. On EMPIAR-10516 and EMPIAR-10345, cryoMV achieved competitive resolution and excelled in generation–reconstruction consistency, motion interpretability, and robustness to particle contami-nation. These results show that leveraging structural priors together with particle evidence reduces unsupported artifacts and improves the reliability of continuous conformations. Although cryoMV currently depends on reference conformations and upstream pose and CTF estimates, future work will focus on reducing these dependencies and extending the approach to more complex heterogeneous data.

